# NOTCH3 signalling controls human trophoblast stem cell expansion and differentiation

**DOI:** 10.1101/2023.07.03.547490

**Authors:** Bianca Dietrich, Kunihs Victoria, Andreas I. Lackner, Gudrun Meinhardt, Bon-Kyoung Koo, Jürgen Pollheimer, Sandra Haider, Martin Knöfler

**Author notes:** co-correspondence: Sandra Haider and Martin Knöfler, Department of Obstetrics and Gynecology, Reproductive Biology Unit, Placental Development Group, Medical University of Vienna, Währinger Gürtel 18-20, 5Q, 1090 Vienna, Austria.

## Abstract

Failures in growth and differentiation of the early human placenta are associated with severe pregnancy disorders such as preeclampsia and fetal growth restriction. However, regulatory mechanisms controlling development of its epithelial cells, the trophoblasts, remain poorly elucidated. Using trophoblast stem cells (TSCs), trophoblast organoids (TB-ORGs) and primary cytotrophoblasts (CTBs) of early pregnancy, we herein show that autocrine NOTCH3 signalling controls human placental expansion and differentiation. NOTCH3 receptor was specifically expressed in proliferative CTB progenitors and its active form, the nuclear NOTCH3 intracellular domain (NOTCH3-ICD), interacted with the transcriptional co-activator Mastermind-like 1 (MAML1). Doxycyclin-inducible expression of dominant-negative MAML1 in TSC lines provoked cell fusion and upregulation of genes specific for multinucleated syncytiotrophoblasts, the differentiated hormone-producing cell type of the placenta. However, progenitor expansion and markers of trophoblast stemness and proliferation were suppressed. Accordingly, inhibition of NOTCH3 signalling diminished growth of TB-ORGs whereas overexpression of NOTCH3-ICD in primary CTBs and TSCs showed opposite effects. In conclusion, the data suggest that canonical NOTCH3 signalling plays a key role in human placental development promoting self-renewal of CTB progenitors.

## INTRODUCTION

Accurate development of the human placenta and its different trophoblast cell types is key to physiological embryonic development and a successful pregnancy outcome. Originating from the trophectoderm the functional units of this unique extraembryonic organ, the placental villi, evolve within 3 weeks post-implantation (Hamilton and Boyd, 1960; Hemberger et al., 2020; James et al., 2012; Knöfler et al., 2019; Turco and Moffett, 2019). The villous structures rapidly expand by branching morphogenesis, generating a surface of 12-14 m^2^ towards the end of gestation, and fulfil all essential tasks of the mature placenta such as fetal nutrition, gas exchange, hormonal adaption of the maternal endocrine system and immunological acceptance of the conceptus (Burton and Fowden, 2015; Erlebacher, 2013; Evain-Brion and Malassine, 2003; Napso et al., 2018). Distinct epithelial trophoblast subtypes that already emerge at the early stages of placental development execute these duties. Extravillous trophoblasts (EVTs) develop in specialized villi, the anchoring villi that attach to the maternal decidua, the endometrium of the pregnant uterus. Upon differentiation, EVTs detach from proliferative cell columns, harbouring NOTCH1/ITGA2^+^ EVT progenitors (Haider et al., 2016; Lee et al., 2018), and invade the decidual stroma and glands ensuring allorecognition and histiotrophic nutrition of the fetus during the early phases of gestation (Burton et al., 2020; Moffett and Shreeve, 2023). EVTs also play a crucial role in controlling adapted blood flow to the placenta by altering uterine vessel function during pregnancy (Burton et al., 2010; Pijnenborg et al., 1983; Pijnenborg et al., 1980). At early stages of development, endovascular EVTs migrate into the maternal spiral arteries and plug the vessels thereby preventing early onset oxygenation and damage of the placenta. However, during establishment of the fetal-maternal connection these EVTs remodel the maternal arteries thereby enlarging their diameter (Pijnenborg et al., 2006). This process allows to precisely regulating blood flow to the developing placenta at later stages of pregnancy when the embryo is feeded by hemotrophic nutrition. In contrast to EVTs that interact with uterine cells (Pollheimer et al., 2018), multinucleated syncytiotrophoblasts (STBs), the second differentiated trophoblast cell type of the placenta, reside in placental floating villi and are surrounded by maternal blood. They develop by cell fusion of underlying cytotrophoblasts (CTBs) representing the proliferative progenitor cell pool of the bi-layered trophoblast epithelium. STBs represent the barrier between maternal and fetal circulation and fulfil a plethora of tasks including secretion of pregnancy hormones, transport of nutrients and waste products as well as oxygen delivery to the growing fetus (Lager and Powell, 2012; Maltepe and Fisher, 2015; Renaud and Jeyarajah, 2022). Defects in the placentation process have been associated with the great obstetrical syndromes including spontaneous abortion, preterm labor, fetal growth restriction (FGR) and preeclampsia (PE) (Brosens et al., 2011). Failures in physiological remodelling of the spiral arteries have been noticed in these disorders provoking malperfusion and subsequent oxidative stress of the placenta (Khong et al., 1986; Pijnenborg et al., 1991; Romero et al., 2011). Abnormal development, expansion and differentiation of trophoblasts could represent underlying causes since CTBs of FGR and PE tissues show alterations in proliferation, STB formation or EVT differentiation (Farah et al., 2020; Lim et al., 1997; Redline and Patterson, 1995; Zhou et al., 1997).

While our knowledge on placental defects in gestational diseases is largely based on histopathological examinations, adequate trophoblast systems allowing in depth investigations of the underlying molecular mechanisms have been poorly developed. In particular, ethical constraints, the limited access to early placental material and the lack of self-renewing trophoblast stem cell models in the past have been a hindrance. However, with the recent establishment of 2-dimensional (2D) human trophoblast stem cells (TSCs) and 3D trophoblast organoids (TB-ORGs) considerably progress has been made (Haider et al., 2018; Okae et al., 2018; Turco et al., 2018). Using these models, developmental signalling pathways, mediated through epidermal growth factor (EGF), Wingless (WNT)-dependent T-cell factors (TCFs), and transforming growth factor β (TGF-β)-activated-SMAD3, have been delineated as crucial regulators of TSC renewal and/or differentiation (Haider et al., 2022; Haider et al., 2018; Okae et al., 2018; Turco et al., 2018). The HIPPO downstream factors TEAD4, YAP and TAZ are also critical for expansion of TSCs and TB-ORGs and abnormal expression of these factors was noticed in different pregnancy complications (Meinhardt et al., 2020; Ray et al., 2022; Saha et al., 2020).

NOTCH represents another signalling cascade that was shown to play a major role in human placentation and trophoblast development (Dietrich et al., 2022; Haider et al., 2017). In the canonical pathway, a membrane-bound ligand of the Serrate-like (JAG1, JAG2) or Delta-like family (DLL1, 3 or 4) interacts with one of the four NOTCH receptors (Kopan and Ilagan, 2009). Subsequently, two sequential proteolytic cleavage steps result in the generation of the transcriptionally active domain of NOTCH (Fortini, 2009; Siebel and Lendahl, 2017; Wolfe and Kopan, 2004). In the first cleavage step, ADAM protease produces the NOTCH extracellular truncation (NEXT). NEXT is then chopped by γ-secretase at the membrane, or in the cytoplasm after endocytosis thereby generating the NOTCH intracellular domain (NOTCH-ICD). The latter translocates into the nucleus with the help of importin α proteins and converts the repressor protein recombination signal binding protein for immunoglobulin kappa J (RBPJκ) into a transcriptional activator by recruiting Mastermind-like (MAML) proteins and other co-activators (Huenniger et al., 2010; Kitagawa, 2016; McElhinny et al., 2008; Siebel and Lendahl, 2017). Repressors of the Hairy/Enhancer of split (HES) and Hey (HES-related with YRPW motif) family members are well known targets genes of NOTCH signalling and control diverse developmental processes (Borggrefe and Oswald, 2009; Siebel and Lendahl, 2017; Weber et al., 2014).

NOTCH regulates trophoblast invasion and the expression of its receptors and ligands dynamically changes during EVT differentiation (Haider et al., 2014; Hunkapiller et al., 2011). NOTCH1 is specifically expressed in the proximal cell column of anchoring villi and has been delineated as a key regulator of EVT development, whereas NOTCH2 was shown to control EVT migration and spiral artery remodelling (Haider et al., 2016; Hunkapiller et al., 2011; Plessl et al., 2015). While first insights into the role of NOTCH in EVT differentiation were unraveled, expression patterns of NOTCH components in early floating villi and their roles in trophoblast stem cell maintenance and STB formation have not been elucidated. Using primary CTBs, TSCs and TB-ORGs we herein demonstrate that canonical NOTCH3 signalling plays a crucial role in trophoblast self-renewal. NOTCH3 is specifically expressed in CTB progenitors and NOTCH3-ICD, binding to its co-activator MAML1, promotes expression of cell cycle and trophoblast stemness genes. Consequently, NOTCH3-ICD fosters expansion of 2D TSCs and 3D TB-ORGs and inhibits differentiation into multinuclear STBs.

## RESULTS

### NOTCH3 and its coactivator MAML1 are specifically expressed in cytrophoblast progenitors of placental tissues and TB-ORGs

Tissue and cellular distribution of NOTCH receptors, their membrane-bound ligands as well as MAMLs were analyzed in isolated first trimester trophoblasts and early placental tissues (Fig.1). To obtain cell pools enriched with villous CTBs and STBs, respectively, a novel two-step protease digestion protocol was developed. After mechanical dissection and removal of villous tips from single placental tissues, the residual material was digested twice allowing sequential isolation of STBs and CTBs (Fig. S1A). Real-time qPCR analyses for makers of trophoblast stemness and cell fusion indicated high purity of the isolated trophoblast subtypes (Fig. S1B). Using these samples abundant mRNA levels of *NOTCH3*, *MAML1-3*, *JAG1* and *DLL1* as well as low levels of *JAG2* and *DLL4* were detected in CTBs and STBs, whereas *NOTCH4* and *DLL3* were absent (Fig. 1A,B). While mRNA levels of the aforementioned genes were present in both villous trophoblast subtypes, protein expression of NOTCH3, MAMLs and ligands was highly restricted. NOTCH3, MAML1 and 3, JAG1 and DLL1 were exclusively detected in isolated CTBs, expressing the stemness markers TEAD4 and YAP1, whereas MAML2 was present in ENDOU/GDF15^+^ STBs (Fig. 1C,D). The 90 kD NOTCH3-NEXT domain was the predominant signal obtained in the Western blot analyses suggesting canonical activation of the pathway. Immunofluorescence of 6^th^ and 12^th^ week placental tissues also indicated specific expression of NOTCH3 in TEAD4^+^ CTBs (Fig. 1E). NOTCH3 immunofluorescence signals were mainly detected in the cytoplasm of CTBs, however, a small fraction (below 1%) of the progenitors showed nuclear NOTCH3-ICD. MAML1 and 3 localized to CTB nuclei as well, while MAML2 was mainly detected in STB nuclei (Fig 1E). NOTCH1 and NOTCH2 were absent from both epithelial trophoblast layers and could only be observed in stromal cells of the villous core (Fig. S2A). Expression in self-renewing TB-ORGs, established from first trimester CTB progenitors, mimicked the patterns monitored in early placental tissues, however with subtle differences (Fig. S2B). MAML1 localized to the outer CTB layer(s) and MAML2 was mainly detected in the centre of 3D TB-ORGs, where syndecan (SDC1)^+^ STBs are formed. However, MAML3 was uniformly expressed. Likewise, STBs in proximity to the TEAD4^+^ CTB layer maintained NOTCH3 expression and only the inner core was negative (Fig. S2B).

**Fig. 1.**
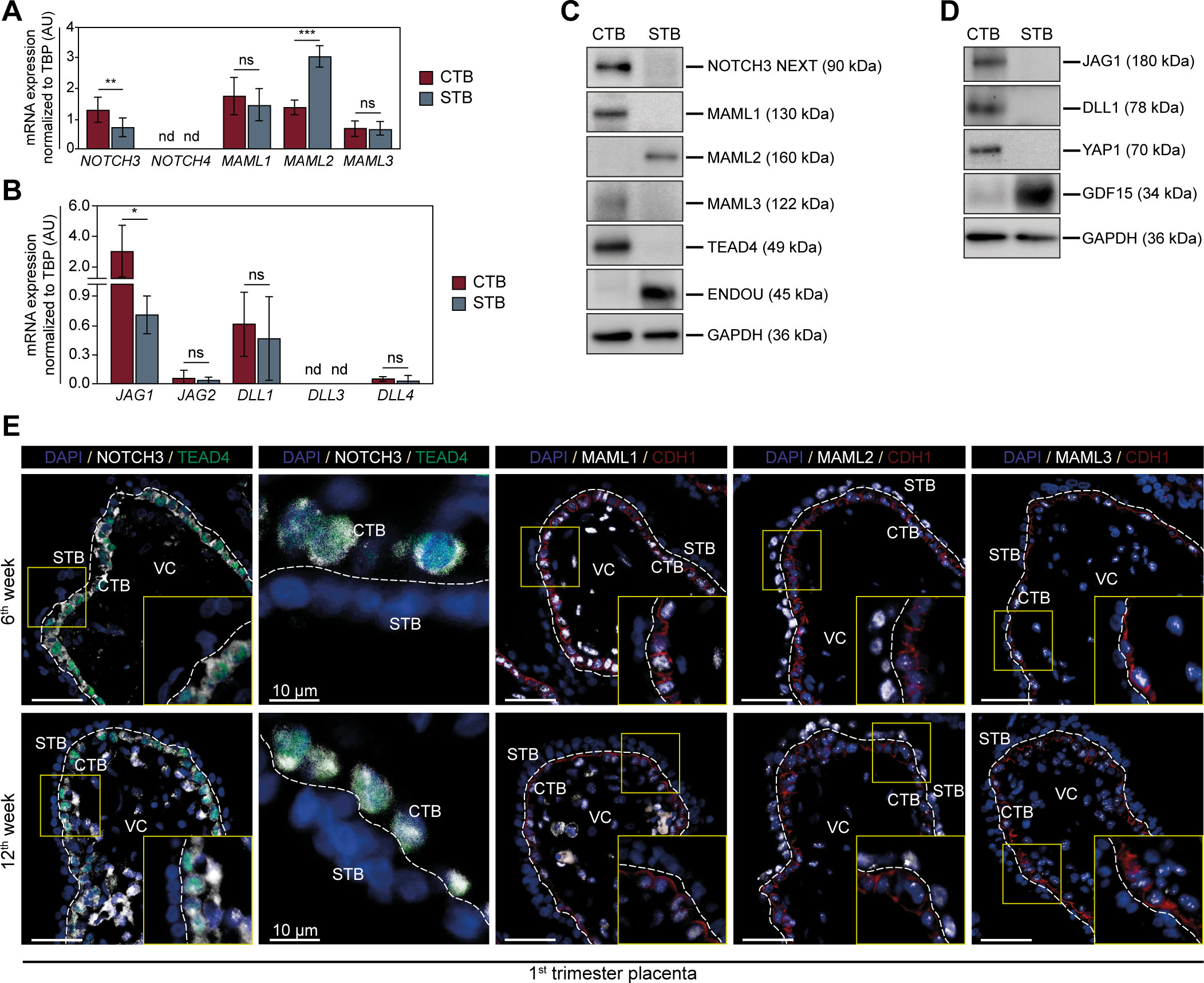
Expression and localization of NOTCH receptors, ligands and MAML co-activators in purified trophoblast subtypes and first trimester placental tissues. (A-B) Relative mRNA expression in matched CTBs and STBs, purified by a two-step protease digestion protocol from first trimester placentae (n=7, 6^th^ to 12^th^ week), was measured by RT-qPCR (duplicates). Data were normalized to transcript levels of TATA box binding protein (*TBP*). Graphs represent mean values±s.e.m. Asterisks denote statistical differences (*P<0.05, **P<0.01, ***P<0.001) as determined by unpaired, two-tailed Student’s *t*-test. ns, not significant. nd, not detectable. AU, arbitrary units; (C-D) Representative Western blots showing protein expression in the isolated trophoblast samples of early placentae (n=3, 7^th^ to 9^th^ week). GAPDH was used as a loading control. (E) In situ localization in first trimester placental tissues. Representative immunofluorescence images of placental sections from 6^th^ (n=3) and 12^th^ week (n=3) of gestation are shown. Stippled line indicates the border between villous cytotrophoblast (CTB) and syncytiotrophoblast (STB). Higher magnifications of inset pictures are shown in the bottom right corner (digitally zoomed). Scale bar: 50 µm. TEAD4 and CDH1 (E-cadherin) mark the CTB cell layer. Nuclei are stained with DAPI. Images with 10 µm scale bar depict selected CTB nuclei expressing NOTCH3-ICD. VC, villous core;

### NOTCH3-ICD interacts with MAML1 in self-renewing TSCs

Self-renewing and fused TSCs showed a similar expression pattern of *NOTCH* receptors and *MAML* co-activators as purified CTBs and STBs, respectively (Fig. 2A). However, these cells expressed low levels of *DLL1* and *NOTCH1* (Fig 2A,B). In accordance with the distribution in primary trophoblast subtypes, NOTCH1 and 3, MAML1 and 3 as well as JAG1 decreased during cAMP-mediated TSC fusion, while MAML2 increased (Fig. 2C-E). Besides the NEXT domain, NOTCH3-ICD was also detectable in protein lysates, which might indicate a high activity of the pathway in proliferating TSCs. (Haider et al., 2016; Rand et al., 2000). Accordingly, immunofluorescence in self-renewing TSCs not only suggested membrane and cytoplasmic localization of NOTCH3, but also a higher proportion of NOTCH3^+^ nuclei compared to primary tissues (Fig. 2E). MAML1 and 3 also localized to nuclei of expanding TSCs, while NOTCH1 was only detected in a small subset of these cells. Immunoprecipitation (IP) using cell lysates prepared from TSCs revealed binding of NOTCH3-ICD to its co-activator MAML1 (Fig. 2F).

**Fig. 2.**
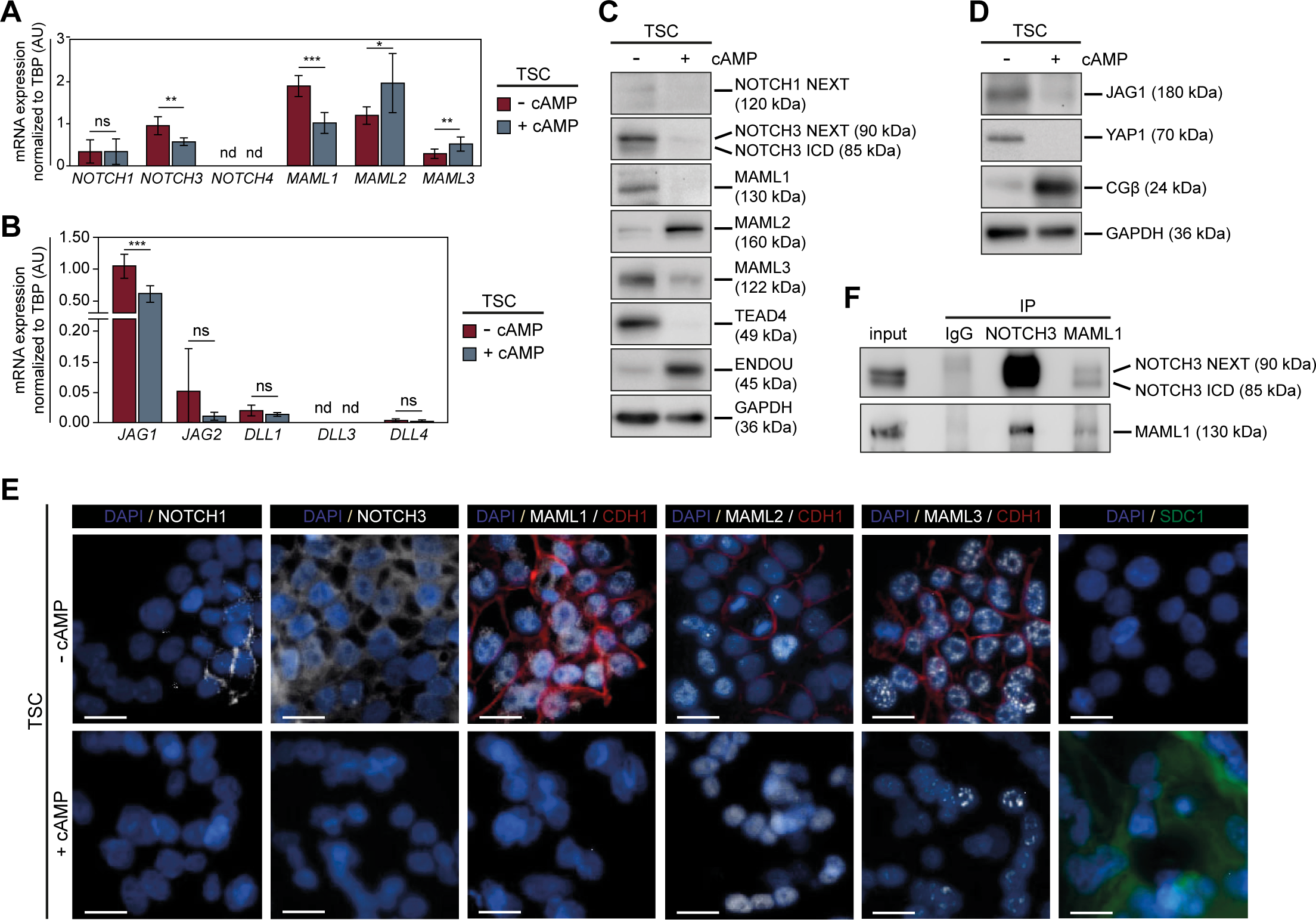
Expression and localization of NOTCH receptors, ligands and MAML co-activators in self-renewing and differentiated trophoblast stem cells. (A-B) RT-qPCR analyses measuring transcript levels in proliferating TSCs (-cAMP; n=6) and differentiated TSCs, treated with forskolin for 5 days (+cAMP, n=6). Data were normalized to *TBP* (AU, arbitrary units). Graphs represent mean values±s.e.m. Asterisks denote statistical differences (*P<0.05, **P<0.01, ***P<0.001) as calculated by unpaired, two-tailed Student’s *t*-test. ns, not significant. nd, not detectable. (C-D) Representative Western blots showing protein expression in proliferating and cAMP-differentiated TSCs (n=3). TEAD4/YAP1 and ENDOU were selected as markers of stemness and cell fusion, respectively. GAPDH was used as loading control. (E) Immunofluorescence in self-renewing and cAMP-treated TSCs (n=3). Scale bar: 25 µM. CDH1 (E-cadherin) and SDC1 were used as markers of expanding and fused TSCs, respectively. (F) Immunoprecipitation in cell lysates, prepared from proliferating TSCs (n=3), using antibodies binding NOTCH3 and MAML1, respectively. A representative example is shown. IgG was used as negative control. IP, immunoprecipitation;

### Bulk RNA-seq of TSCs, expressing dominant-negative MAML1, revealed changes in trophoblast expansion and cell fusion

To specifically inactivate canonical NOTCH3 signalling in self-renewing TSCs, progenitor cells, isolated from three different first trimester placentae, were transfected with a doxycycline (Dox)-inducible dominant-negative (DN) MAML1 construct. In this plasmid the basic domain (BD) of MAML1, required for binding to NOTCH-ICD, was fused to eGFP (Fig. 3A). Thus, truncated DN-MAML1 lacks protein sequences downstream of BD that are required for recruitment of other transcriptional co-activators such p300/CBP, YAP/TAZ or β-catenin (McElhinny et al., 2008; Zema et al., 2020). Three different TSC clones were obtained that expressed fluorescent DN-MAML1 and DN-MAML1 transcripts after induction with Dox (Fig. S3A). Since increased cell fusion was observed in these cultures, we performed kinetic analyses to determine the optimal time span for Dox treatment. Transcript levels of trophoblast progenitor markers where lowest after six days of Dox addition, while STB markers were highest (Fig. S3B). This condition was selected for all subsequent experiments using DN-MAML1-expressing TSC lines. Bulk RNA-seq and bioinformatics analyses of the three clones revealed 1182 upregulated and 637 downregulated genes after Dox induction (Fig. 3B). Well known STB markers such as genes encoding chorionic gonadotrophin β (*CGB3*, *CGB5*, *CGB7*, *CGB8*), the syncytins (*ERVW1*, *ERVFRD1*), *SDC1*, leptin (*LEP*) and growth and differentiation factor 15 (*GDF15*) were among the upregulated genes. The full list of regulated genes is available at GEO (accession number GSE233140). In contrast, genes for trophoblast stemness (*TEAD4*, *MYC*, *LGALS1*) and proliferation/mitosis (*CCNA2*, *CCNB1* and *2*, *CCND1*, *CCNE2*, *CDK1*) were significantly decreased. Although the three TSC lines differed in the PCA (Fig. S3C), relative up- or downregulation of these marker genes was consistent in each of the cultures (Fig. 3C).

**Fig. 3.**
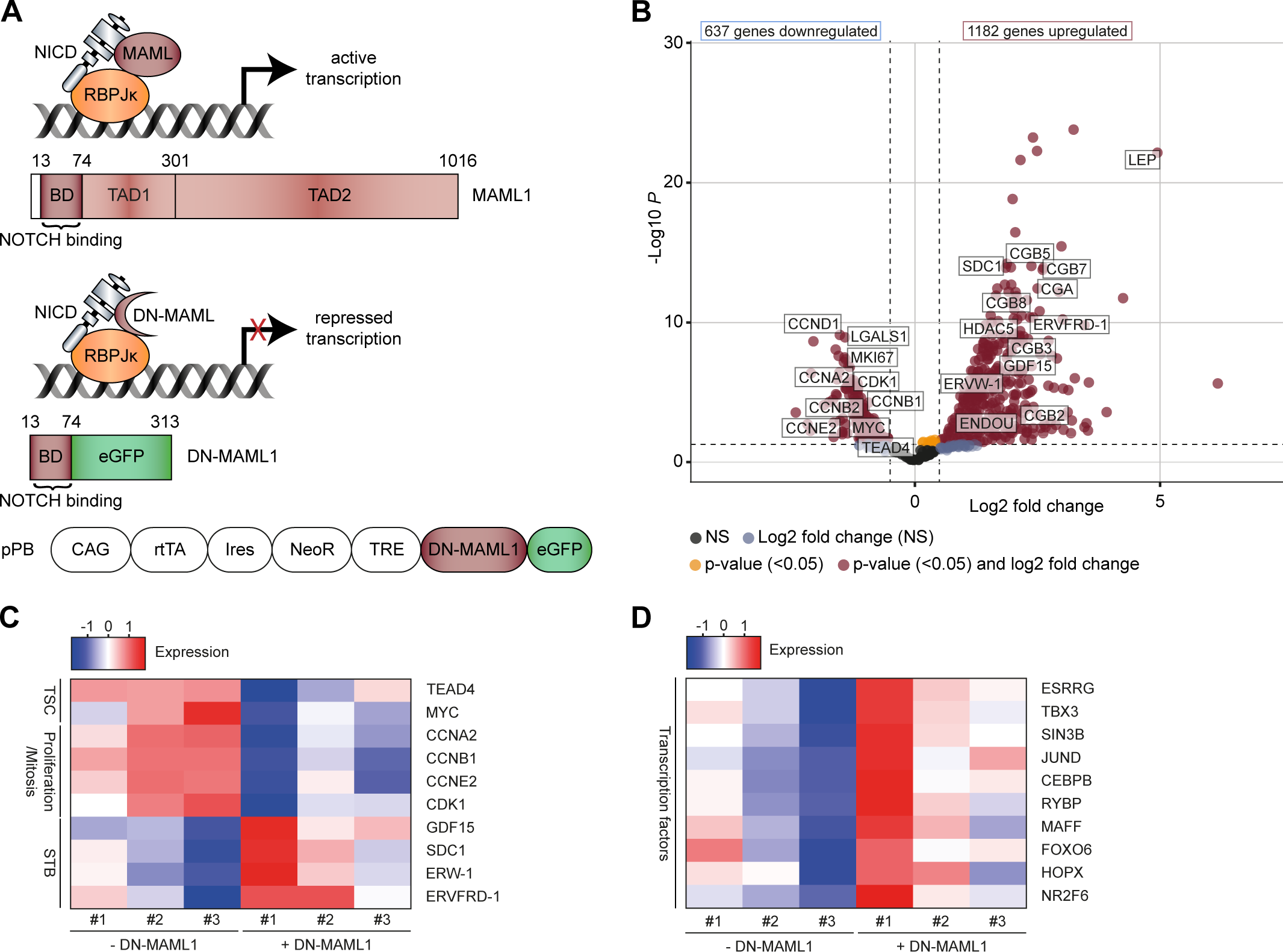
Genome-wide expression analyses of TSCs clones expressing dominant-negative (DN) MAML1. (A) Schematic representation of transcriptionally active NOTCH(N)-ICD-MAML-RBPJk versus an inactivated complex binding DN-MAML1 that lacks transcription activation domain 1 and 2 (TAD1 and 2). BD, basic domain. The graph at the bottom illustrates the structure of the piggyBac (pPB) vector expressing DN-MAML1, eGFP, and the reverse Tet-repressor (rtTA) for Dox-inducible expression. (B) Volcano blot showing genome-wide gene expression, six days after expression of DN-MAML1 in the three different TSCs clones. Among the statistically regulated mRNAs, specific markers of expanding CTBs and differentiated STBs are highlighted. Dots depict individual transcripts, colored according to P values and log2 fold change (DESeq2, standard parameters). NS, not significant; (C) Heatmap showing selected markers of TSCs, proliferation/mitosis and STBs in the three individual TSC clones in the absence or presence of DN-MAML for six days. (D) Heatmap depicting a previously proposed transcription factor network associated with STB differentiation in the DN-MAML-expressing TSC clones with and without Dox treatment.

To study possible side effects of the Dox supplementation we also profiled non-transfected TSCs after six days of incubation with the antibiotic. Treated and untreated TSCs were highly similar in the PCA and none of aforementioned trophoblast progenitor markers and STB-specific genes were significantly changed (Fig. S3C,D). Only two of the 21 genes, significantly upregulated by Dox, were also increased upon DN-MAML1 expression, whereas none of the eight Dox-downregulated transcripts was among the DN-MAM1-suppressed genes (Fig. S3E). DN-MAML1 induction also increased transcription factors that were previously defined as a network associated with STB differentiation (Chen et al., 2022) in the three TSC clones (Fig. 3 D).

### Inducible expression of dominant-negative MAML1 in TSCs suppresses markers of trophoblast stemness and upregulates STB-specifc genes

To verify bulk RNA-seq data, trophoblast subtype-specific markers were analyzed in the three different TSC lines (Fig. 4). Upon DN-MAML1 expression mRNA and/or protein levels of trophoblast stemness (TEAD4, YAP1, MYC) and proliferation/mitosis-associated genes (cyclin A, cyclin B, cyclin D) were diminished while STB-specific genes (GDF15, *ENDOU*, CGβ) were increased (Fig. 4A,B). DN-MAML1 expression also downregulated endogenous levels of the NOTCH3-NEXT and NOTCH3-ICD in the different TSC clones. Of the relevant HES and HEY family members only *HEYL* was suppressed by DN-MAML1, while *HEY1* was upregulated in two of the three clones (Fig. S4). *HES1* and *HES7* differed between the TSC lines, while *HES5* and *HEY2* were absent.

**Fig. 4.**
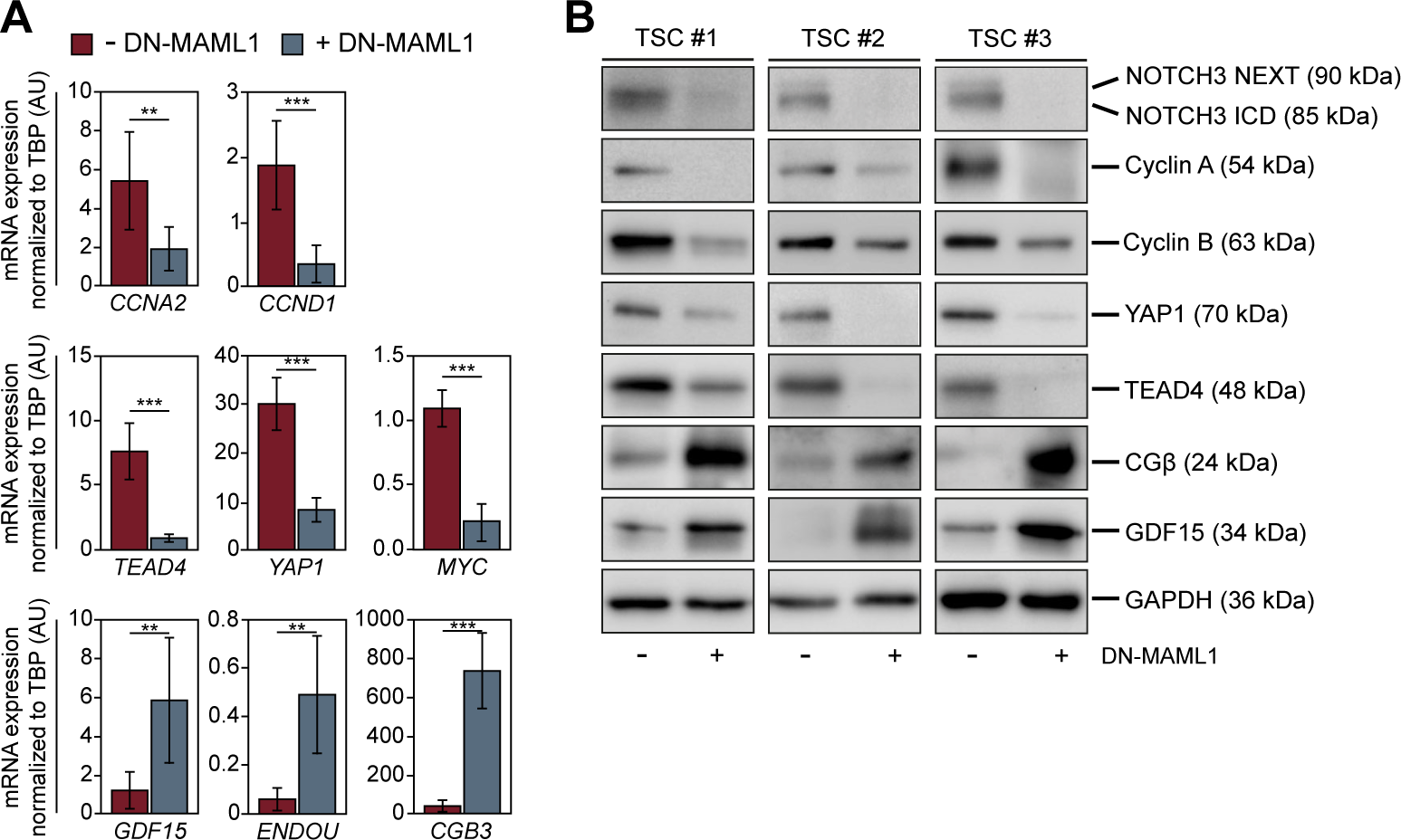
Inducible expression of dominant-negative (DN) MAML1 affects markers of trophoblast stemness, proliferation and cell fusion. (A) Real-time qPCR measuring mRNA expression (n=6, the three clones in duplicates) in untreated and Dox-treated (six days) TSCs. Graphs represent mean values±s.e.m. Asterisks denote statistical differences (**P<0.01, ***P<0.001) as determined by unpaired, two-tailed Student’s *t*-test. (B) Representative Western blots (n=3 per TSC line) showing changes in the protein expression of selected trophoblast markers with and without DN-MAML1 expression in the three different TSC clones. GAPDH was used as loading control.

### NOTCH3 inhibition decreases proliferation and elevates cell fusion in TSCs and TB-ORGs

Next, we investigated the biological effects of DN-MAML on TSC expansion and STB formation. Dox-dependent expression of the inhibitory protein provoked downregulation of proliferation, measured by EdU labelling, while cell fusion, analyzed by the formation of the SDC1^+^ area, increased in the three TSC lines (Fig. 5A,B). Notably, the three clones differed in proliferation and particularly in their capacity for STB formation upon DN-MAML1 expression (Fig. 5A,B). In agreement, chemical blockage of the pathway with the γ-secretase L-685,458 inhibitor impaired growth of self-renewing TB-ORGs (Fig. 5C). L-685,458 also decreased cyclin A and YAP1, but and increased CGβ expression in the 3D system (Fig. 5D). Another inhibitor of NOTCH cleavage, DBZ, showed similar results (Fig. S5A,B).

**Fig. 5.**
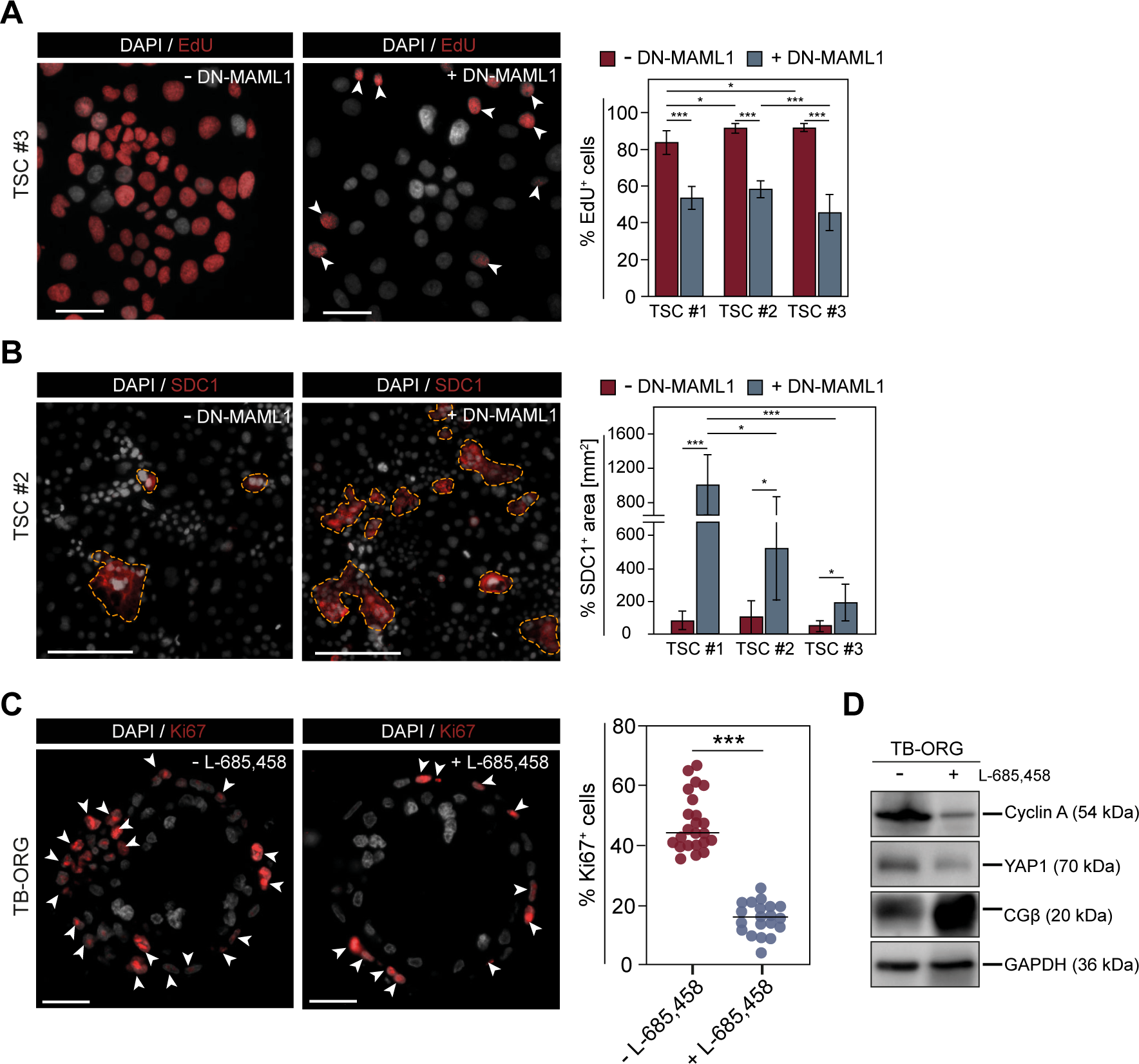
Genetic and chemical inhibition of NOTCH downregulates proliferation and increases cell fusion in self-renewing TSCs and TB-ORGs. (A) EdU labelling of DN-MAML1 transfected TSCs in the absence or presence of Dox for 6 days. Representative immunofluorescence images of TSC line 3 are depicted. Scale bar: 50 µM. To determine the EdU^+^/DAPI ratio of nuclei, for each TSC line and condition 10 individual areas each containing between 550 and 700 nuclei were manually counted. Graphs represent mean values±s.e.m. Asterisks denote statistical differences (*P<0.05, ***P<0.001) as determined by ordinary one-way ANOVA and two-tailed Turkey post-test. (B) SDC1^+^ areas in TSCs with and without DN-MAML1 expression. Representative immunofluorescence pictures (TSC line 2) with multinucleated SDC1^+^ areas, encircled by stippled lines, are shown. Scale bar: 200 µM. To measure the extent of SDC1^+^ areas, 10 individual pictures, each harboring between 720 and 870 nuclei, were analyzed for each TSC line and condition. Graphs represent mean values±s.e.m. Asterisks denote statistical differences (*P<0.05, ***P<0.001) as determined by ordinary one-way ANOVA and two-tailed Turkey post-test. (C) Immunofluorescence in tissue sections of TB-ORGs treated with the γ-secretase inhibitor L-685,458. Representative pictures of TB-ORGs derived from a 6^th^ week primary CTB preparation are shown. Scale bar: 25 µM. For evaluation of the number of Ki67^+^ nuclei, 23 untreated and 20 L-685,458-treated TB-ORGs derived from two different first trimester placentae (6^th^ and 7^th^ week) were analyzed. Graph represents mean values±s.e.m. Asterisks denote statistical differences (***P<0.001) as determined by unpaired, two-tailed Student’s *t*-test. (D) Representative Western blot showing protein expression in TB-ORGs (n=3) supplemented with L-685,458. GAPDH was used as loading control.

### Overexpression of NOTCH3-ICD promotes stemness and proliferation in primary trophoblast progenitors and TSCs

To assess the role of transcriptionally active NOTCH3 in trophoblast self-renewal, trophoblast progenitors were transiently transfected with a plasmid encoding NOTCH3-ICD (Fig. 6). Primary CTBs and TSCs overexpressing NOTCH3-ICD showed elevation of markers for trophoblast stemness (TEAD4, YAP1) and proliferation/mitosis (cyclin A, cyclin B, cyclin D), while CGβ was downregulated (Fig. 6A,D). Accordingly, NOTCH3-ICD increased the number of EdU^+^ cell in TSC cultures, whereas the multinuclear SDC1^+^ area was diminished (Fig. 6B,C). Moreover, NOTCH3-ICD upregulated endogenous NOTCH3-NEXT signals in the two cell types (Fig. 6A,D). Compared to controls, the active domain also increased expression of the full-length receptor in primary CTBs and TSCs (Fig. S6A,B). In agreement with that, Dox-induction of DN-MAML1 in the TSC lines decreased the faint signals of 270 kD NOTCH3 (Fig. S6C).

**Fig. 6.**
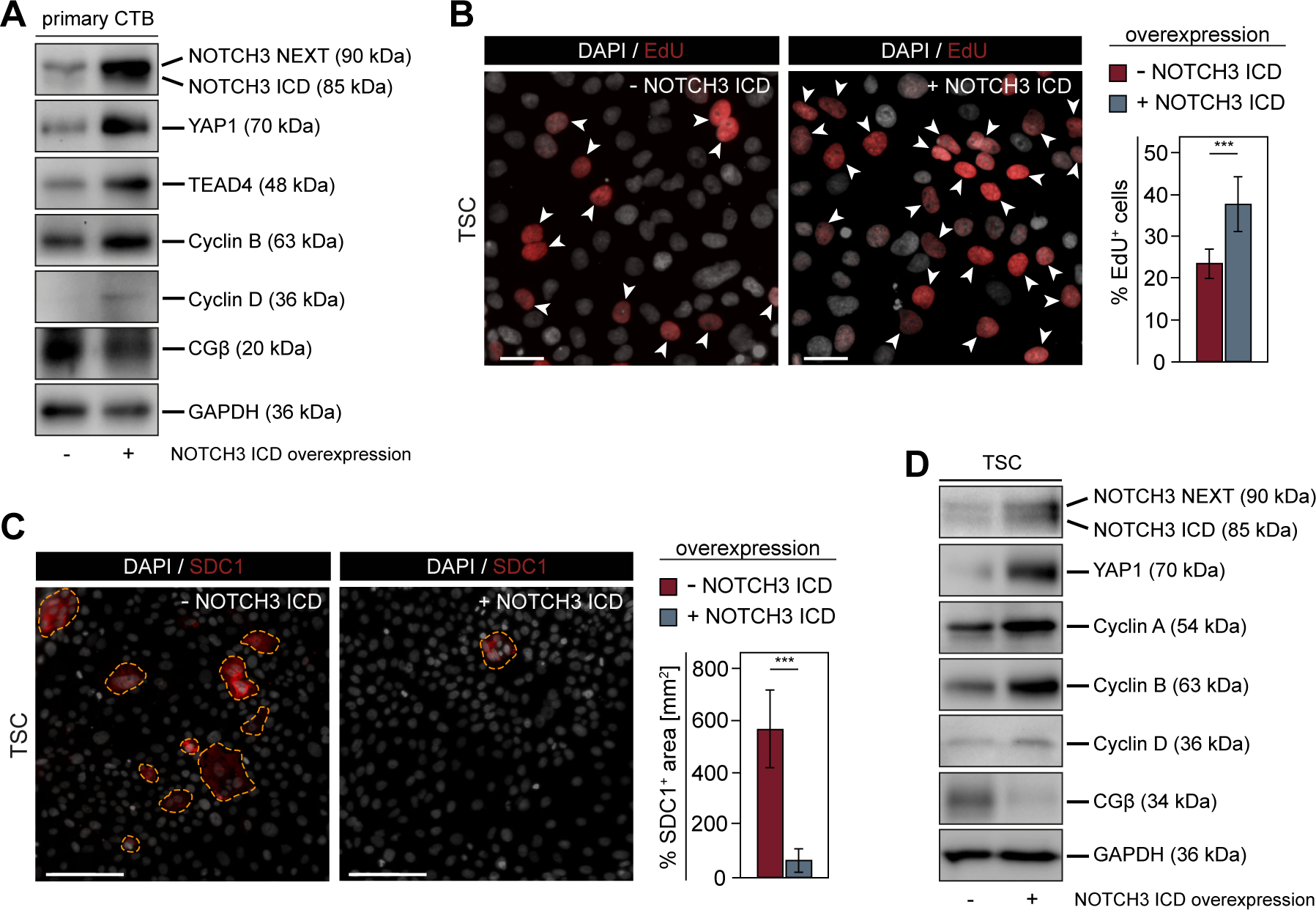
NOTCH3-ICD promotes trophoblast expansion and impairs differentiation. Representative Western blots detecting protein expression of trophoblast stemness- and proliferation/mitosis-associated genes upon overexpression of NOTCH3-ICD in (A) primary CTBs (n=3) and (D) TSCs (n=3). GAPDH was used as loading control. (B) Representative immunofluorescence images showing EdU labelling in TSCs that express NOTCH3-ICD. Scale bar: 50 µM. Percentage of EdU^+^ TSCs (n=9) was measured by counting EdU^+^/DAPI ratio of 10 individual areas per experiment and condition, each containing between 800 and 950 nuclei. Graphs represent mean values±s.e.m. Asterisks indicate statistical differences (***P<0.001) as determined by unpaired, two-tailed Student’s t-test. (C) Immunofluorescence pictures illustrating SDC1^+^ areas (encircled by stippled lines) in the absence or presence of exogenous NOTCH3-ICD. A representative experiment is shown. Scale bar: 200 µM. To determine the size of SDC1^+^ areas, 10 different regions of NOTCH3-ICD-expressing TSCs (n=9) and controls, each containing between 650 and 800 nuclei, were evaluated. Graphs represent mean values±s.e.m. Asterisks denote statistical differences (***P<0.001) as determined by unpaired, two-tailed Student’s t-test.

### Discussion

Recent omics approaches have been increasing our knowledge about human placental and trophoblast development. For example, single cell analyses of primary tissues, TSCs and TB-ORGs gave novel insights into underlying mechanisms, the diversity of trophoblasts and their putative interactions with other placental and decidual cell types (Arutyunyan et al., 2023; Liu et al., 2018; Shannon et al., 2022; Vento-Tormo et al., 2018). Chromatin immunoprecipitation and sequencing has allowed to establishing a landscape of TSC enhancers that harbour binding sites for pivotal transcriptions factor of trophoblast development (Frost et al., 2023; Kim et al., 2023). Moreover, integrated bioinformatics of diverse data sets have suggested signalling molecules and networks that likely play major roles in trophoblast self-renewal (Chen et al., 2022; Cox and Naismith, 2022). While functional analyses that underpin the molecular roles of most of these genes are missing, various transcriptional regulators have been studied in detail. For instance, HIF, ASCL2, GCM1, and TCF4 have been identified as crucial regulators of EVT differentiation (Jeyarajah et al., 2022; Meinhardt et al., 2014; Varberg et al., 2021; Wakeland et al., 2017), whereas factors such as TEAD4, GATA2, MSX2 and TFAPC promote TSC self-renewal (Chen et al., 2022; Hornbachner et al., 2021; Kim et al., 2023; Saha et al., 2020). Transcriptional co-activators, i.e. enhancer-associated p300, and the HIPPO factors TAZ and YAP, are also critical for trophoblast stemness, the latter actively repressing genes associated with cell fusion (Kim et al., 2023; Meinhardt et al., 2020; Ray et al., 2022).

In the present paper, we show that another transcriptional regulator, the NOTCH3-ICD plays a fundamental role in TSC expansion and differentiation. NOTCH3 is the only receptor expressed by villous CTB progenitors of first trimester placental tissues and self-renewing CTBs in TB-ORGs, while TSCs additionally display low levels of NOTCH1 (Fig.1 and 2). NOTCH1 was not uniformly expressed and appeared in small cell clusters of expanding TSCs. Notably, placental NOTCH1 expression was previously identified as a marker of proliferative cell column trophoblasts representing the EVT progenitor pool of anchoring villi (Haider et al., 2016). NOTCH1-ICD has been delineated as a crucial regulator of EVT progenitor formation and survival, which represses the villous CTB progenitor phenotype by downregulating the self-renewal markers TEAD4 and p63, and activates proximal cell column-specific genes (Haider et al., 2016). However, NOTCH1 is absent from the bi-layered trophoblast epithelium of first trimester placental villi and can only be detected in the underlying stroma (Haider et al., 2016) and (Fig. S2A). Hence, upregulation of NOTCH1 could be beneficial for the viability of TSCs upon cultivation in 2D. On the other hand, passaging of TSCs in 2D could allow for the expansion of some of the EVT progenitors, while 3D TB-ORG formation might suppress their abundance. Indeed, TB-ORGs established from the first established TSC lines showed higher NOTCH1 expression than TB-ORGs isolated from freshly prepared patient material (Sheridan et al., 2021).

Akin to EVT differentiation (Haider et al., 2014; Hunkapiller et al., 2011), NOTCH, ligands and MAMLs are dynamically regulated during STB formation. NOTCH3, JAG1, DLL1 and MAML1 and 3 are specific CTB progenitor markers of early placental tissues suggesting that these factors could be associated with trophoblast self-renewal (Fig. 1). Protein levels of these factors were absent from STBs and decreased during c-AMP mediated cell fusion of TSCs (Fig. 2). Downregulation of NOTCH3 was also observed during in vitro EVT differentiation of primary CTBs upon cultivation on fibronectin, reinforcing the idea that the particular receptor is crucial for trophoblast proliferation (Haider et al., 2014). Differently to MAML1 and 3, MAML2 was upregulated during cell fusion. In the absence of any NOTCH receptor, MAML2 likely fulfils a differential role in STBs. Indeed, MAML proteins affect nuclear functions of other developmental regulators in different tissues (Zema et al., 2020). Notably, activation of these factors might predominantly occur when NOTCH is inactive, suggesting that NOTCH-ICDs are the preferred binding partners of MAMLs (McElhinny et al., 2008; Shen et al., 2006). Whereas protein expression of NOTCH3, JAG1 and DLL1 was strongly regulated during cell fusion in situ and in vitro, their transcript levels showed modest changes. Indeed, NOTCH receptors, ICDs and ligands are subject to different posttranslational modifications, intracellular trafficking routes, and degradation pathways to fine-tune the signal output (Kopan and Ilagan, 2009; Siebel and Lendahl, 2017). Hence, NOTCH signalling in trophoblasts may not require strict regulation at the mRNA level.

In situ, NOTCH3 was detected at the membrane and in the cytoplasm suggesting basal activation of the canonical pathway by ADAM cleavage and internalization of the NOTCH3-NEXT fragment as occurs in other cellular systems (Siebel and Lendahl, 2017). Accordingly, NOTCH3-NEXT was abundant in protein lysates of primary CTBs (Fig. 1). The transcriptional co-activator NOTCH3-ICD, produced by γ-secretase in the subsequent proteolytic step, was noticed in a small percentage of CTB progenitors of early placental tissues, but was hardly detectable in cellular extracts. However, NOTCH3-ICD was present in lysates of TSCs and a higher proportion of these cells showed nuclear staining compared to primary CTBs (Fig. 2.).

We assume that variances in the proliferation rates of in situ CTB progenitors versus in vitro-cultivated TSCs could provide an explanation for the observed differences. TSCs, expanding in the presence of EGF, the WNT activator CHIR99021, and the TGF-β inhibitor A8301, likely exhibit elevated growth rates. This might also explain the subtle alterations of NOTCH3 regulation in TB-ORGs, undergoing self-renewal in the presence of the same factors. In agreement with the staining pattern in early placentae, NOTCH3 specifically localized to CTB progenitors of TB-ORGs (Fig. S2B). However, in contrast to its restricted in situ expression, NOTCH3 downregulation was delayed during spontaneous STB formation in this system. Nonetheless, detection of NOTCH3-ICD in situ, which only has a half-life of about 45 min in the presence of MAML (Fryer et al., 2004; Hein et al., 2015), and the functional assays discussed below strongly suggest a role of canonical NOTCH3 signalling in self-renewal of CTB progenitors. The way NOTCH3 is activated in these cells and whether NOTCH ligands are truly involved remains subject to future investigations. Classically, activation of the NOTCH pathway requires juxtaposed cells, expressing receptor and ligand, respectively, whereas cis-expression is thought to elicit inhibitory signals (D’Souza et al., 2008; Siebel and Lendahl, 2017). Yet, trans-expression could not be observed in early placental tissues since NOTCH receptors and ligands co-localize in the different trophoblast subtypes (Haider et al., 2014; Hunkapiller et al., 2011). However, among all NOTCH receptors, NOTCH3 is most easily cleaved (Choy et al., 2017). NOTCH3 shows basal signalling activity in other cells and its activation can occur in the absence of a ligand (Xu et al., 2015). Moreover, cis-expression with JAG1, which is also the major ligand of TSCs and primary CTBs (Fig. 1 and 2), was shown to promote NOTCH3-dependent gene expression (Pelullo et al., 2014). Overall, these data suggest that the mechanisms of NOTCH3 activation are highly variable and context dependent. To exert their biological roles, ICDs of the different NOTCH receptors displace transcriptional repressors from RBPJk complexes and recruit co-activators such as p300, PCAF, MAML and others (Borggrefe and Oswald, 2009; Siebel and Lendahl, 2017). IP experiments showed that the NOTCH3-ICD binds MAML1 in self-renewing TSCs (Fig. 2), suggesting that functional trimeric protein complexes between RBPJk, NOTCH3-ICD and MAML1 are formed. Indeed, RBPJk and NOTCH-ICD each cannot bind MAML as single proteins, only when a complex is built between the two (Nam et al., 2003). This provided the basis for the development of the pan-NOTCH inhibitor DN-MAML that allowed investigating the role of canonical NOTCH signalling in many different biological contexts (Siebel and Lendahl, 2017). Since DN-MAML lacks TAD1 and 2, required for binding of other regulators, it is thought to act as a specific inhibitor of the NOTCH pathway (McElhinny et al., 2008; Zema et al., 2020). Here, we designed a truncated DN-MAML1 protein, harboring the sequence of the BD fused to GFP, and used a Dox-dependent plasmid for inducible expression in TSCs (Fig. 3). G418-seletion yielded three stable DN-MAML1 expressing TSC lines that were subjected to genome-wide expression analyses. The three clones varied in the PCA in the absence or presence of Dox, which could be explained by distinct origins of the transfected trophoblast cells. Indeed, different clusters of CTB progenitors have been described in placental tissues, primary cultures and TB-ORGs (Arutyunyan et al., 2023; Liu et al., 2018; Shannon et al., 2022; Vento-Tormo et al., 2018). However upon Dox-induced DN-MAML1 expression, markers characteristic for STBs were upregulated in all three TSC lines suggesting that inhibition of basal NOTCH3 activity promotes cell fusion (Fig. 3 and 4). DN-MAML1 elevated activators of cell fusion (*ERVW1*, *ERVFRD1*), a cluster of genes (*CGA*, *CGB3*, *CGB5*, *CGB7*, *CGB8*) encoding the pregnancy hormone CG, other STB-secreted factors (*LEP*, GDF15) as well as a set of transcription factors associated with syncytialization (Fig. 3D). Moreover. *HDAC5*, controlling the activity of the master regulator of trophoblast cell fusion, GCM1, was also upregulated (Chang et al., 2013; Jeyarajah et al., 2022). In contrast, DN-MAML1 suppressed genes of trophoblast stemness (TEAD4, YAP1, *MYC, LGALS1*) as well as regulators of cell cycle progression and mitosis (Cyclin A and B, *CCND1*, *CCNE2*, *CDK1*). Accordingly, Dox-dependent expression of the NOTCH inhibitor decreased TSC expansion by around half, while cell fusion was strongly increased (Fig. 5A and B). Notably, reduction of proliferation in the presence of DN-MAML1 was similar in the three lines, while their capacity for cell fusion differed. Distinct origins of the progenitors could again provide an explanation for the latter. Similar to TSCs, inhibition of γ-secretase diminished proliferation and elevated CGβ production in TB-ORGs, suggesting that the NOTCH pathway is also crucial for TSC expansion in 3D (Fig. 5C and D).

Some of the genes, downregulated by DN-MAML1 have been identified as direct targets of NOTCH/NOTCH3-ICD such as *CCND1*, *CCNE*, and *MYC*, which is also bound by NOTCH1-ICD in EVT progenitors (Borggrefe and Oswald, 2009; Haider et al., 2016; Man et al., 2012; Siebel and Lendahl, 2017; Zender et al., 2016). Interestingly, NOTCH3 also maintains its own expression in TSCs (Fig. 4B), as shown in other cells (Liu et al., 2009; Weng et al., 2006), suggesting that an autocrine NOTCH3 loop governs trophoblast expansion. Furthermore, we speculate that DN-MAML1-induced STB formation and upregulation of fusogenic genes, such as the syncytins, could be largely a consequence of the downregulation of proliferation markers and exit from the cell cycle. Indeed, progression into the G0 phase of the cycle and G0-restricted expression of syncytin 2 was shown to be necessary for developing functional STBs (Lu et al., 2017). To exclude any side effects of DN-MAML1 on other signalling pathways, TSC and CTB progenitors were also transfected with NOTCH3-ICD. Overexpression of the active domain increased TCS expansion, cyclins and trophoblast stemness genes while cell fusion and CGβ expression was suppressed (Fig. 6). Noteworthy, transient expression of NOTCH3-ICD also provoked upregulation of the full-length NOTCH3 receptor and its NEXT domain (Fig.6. and Fig. S6), again suggesting that the basal activity of the pathway is maintained in an autocrine fashion. In summary, the results of the NOTCH3-ICD experiments are in line with observations on the DN-MAML1-expressing TSC lines reinforcing the idea that NOTCH3 plays a major role in trophoblast expansion.

In conclusion, the present data suggest that NOTCH3 is crucial for early placental development (Fig. 7). Villous CTB progenitors and TSCs specifically produce NOTCH3 and its cleavage product NOTCH3-ICD sustaining NOTCH3 expression and, consequently, basal activity of the pathway. NOTCH3-ICD, binding to the co-activator MAML1, maintains expression of cell cycle and stemness genes that are required for trophoblast self-renewal. As a result, the default differentiation pathway of floating placental villi, formation of STBs by cell fusion, is suppressed.

**Fig. 7.**
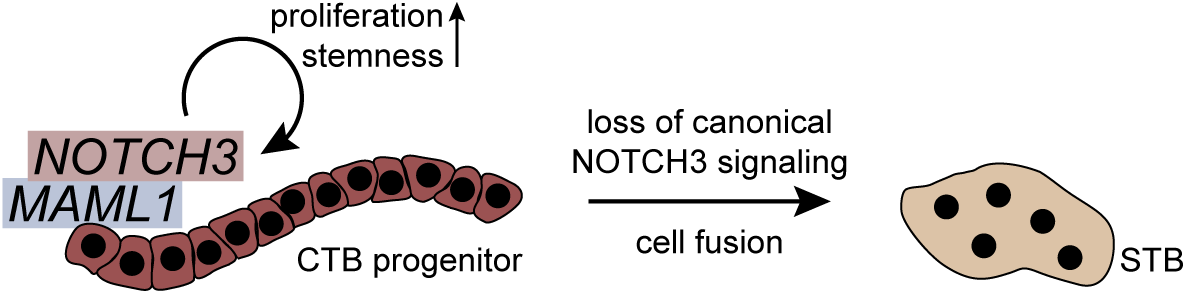
Schematic depiction of the role of canonical NOTCH3 signalling human trophoblast cells. Active NOTCH3-ICD, interacting with the transcriptional co-activator MAML1, promotes self-renewal of CTB progenitors and TSCs and maintains its expression in an autocrine manner, whereas downregulation of the NOTCH3 pathway results in STB formation.

## MATERIALS AND METHODS

### Tissue collection

First-trimester placental tissues (6^th^ to 12^th^ week of gestation) were obtained from elective pregnancy terminations performed by Gynmed Clinic, Vienna. Utilization of tissues and experimental procedures were approved by the ethical committee of the Medical University of Vienna (no. 084/2023) requiring written informed consent from donating women. If not stated otherwise, cell isolations were performed from single placentae.

### Isolation of CTBs and STBs from first trimester placentae using sequential digestions

Placental tissues (6^th^-12^th^ week of gestation) were washed with Mg^2+^/Ca^2+^-free Hanks balanced salt solution (HBSS, Gibco) and villus-trees were mechanically dissected. Then, all villous tips were thoroughly cropped and removed to avoid contamination with cell columns. Two digestion steps were performed using 5 ml digestion solution/ml tissue for 10 and 15 min, respectively, in a 37°C water bath with careful inversion every 2-3 minutes. The digestion solution was composed of HBSS containing 0.25% trypsin (Gibco) and 1,25 mg/ml DNAse I (Sigma). Each digestion reaction was stopped by using 10% FBS ([vol/vol], Sigma), and the remaining tissue was collected and further processed. The cells resulting from both digestions were treated separately: the first digestion solution was enriched for STBs and the second for CTB progenitor cells. Contaminating red blood cells (if present) were removed upon incubation with erythrocyte lysis buffer (155 mM NH_4_Cl, 10 mM KHCO_3_, 0.1 mM EDTA, pH 7.3) for 5 min at room temperature. Afterwards, both cell populations were washed with 1x phosphate buffered saline (PBS; PanBiotech) and stored as cell pellets for subsequent real-time qPCR and Western blotting analyses. In addition, tissue was put aside before and after cropping of the villous tips as well as after each digestion step, fixed and embedded in paraffin. Using a microtome 1.5 µm sections of these tissues were prepared and stained with Hematoxylin & Eosin (H&E) using standard procedures.

### Isolation and Cultivation of Trophoblast Stem Cells from First-Trimester Placentae

Trophoblast stem cells (TSCs) were isolated by three consecutive enzymatic digestion steps followed by density gradient centrifugation according to protocols mentioned elsewhere (Haider et al., 2018). Briefly, placental tissues (6^th^-8^th^ week of gestation) were washed with HBSS, villi were cut and further chopped into small pieces (1-3 mm). Three digestion steps were performed as described above. Whereas the first digestion was discarded, the second and third digestions, mainly containing the trophoblast progenitor cells, were pooled and purified using a Percoll gradient [10-70% (vol/vol)]. Cells were collected between 35% and 50% of Percoll layers, while contaminating red blood cells (if present) were removed by incubation with erythrocyte lysis buffer. Afterwards, cells were washed with HBSS and either seeded onto fibronectin-coated culture dishes using culture medium that promotes trophoblast stemness (TSC medium) or processed for organoid formation as described below. TSC medium contained DMEM/F12 (Gibco) supplemented with 1x B-27 (Gibco), 1x Insulin-Transferrin-Selenium-Ethanolamine (ITS-X; Gibco), 1 µM A83-01 (Tocris), 50 ng/ml recombinant human epidermal growth factor (rhEGF; R&D Systems), 2 µM CHIR99021 (Tocris) and 5µM Y27632 (Santa Cruz). Cells were passaged with 80-90% confluence at a split ratio of 1:2-1:4 using TrypLE (Gibco) for 8 min at 37°C. The trypsinization reaction was stopped with 10% FBS (vol/vol) and cells were pelleted by centrifugation for 5 min at 1500 rpm and then seeded again or subjected to cryopreservation. For cryopreservation, harvested TSCs were re-suspended in Cellbanker2 (Zenoaq) and frozen at -80°C according to the manufacturer’s instructions. Molecular characterization and differentiation of TSCs as described below was performed at P3-P6.

### In vitro cell fusion of trophoblast stem cells

TSCs were grown to approximately 60-70% confluence in TSC medium prior to induction of syncytialization. STB formation was performed as previously published (Okae et al., 2018), however with minor modifications. TSCs were fused after removal of A83-01, rhEGF and CHIR99021 from the TSC medium by adding 2 mM forskolin. Medium was replaced after 3 days and the cells were harvested using TrypLE at day 5 or fixed for immunofluorescence staining. To remove a possible contamination with TSCs that have spontaneously differentiated into HLA-G^+^ EVTs, harvested cells were re-suspended in MACS buffer (MACS Miltenyi Biotec) followed by magnetic cell separation of HLA-G-negative cells using HLA-G-PE antibodies (PE1P-292-C100, Exbio) and anti-PE MicroBeads (MACS Miltenyi Biotec) as instructed by the manufacturer. As fused TSCs are large and would become stuck in the MACS column, negative selection of forskolin-treated cells was performed by using EasySep cell separation magnet (stemcell technologies). For later usage, cell pellets were stored at -80°C.

### Trophoblast Organoid Formation and Cultivation

Trophoblast organoids (TB-ORGs) were established and cultivated as previously published (Haider et al., 2018) with minor modifications. Either freshly isolated trophoblast progenitor cells or cultured TSCs (P3-P6) were used for formation of TB-ORGs. Briefly, cells were washed in ice-cold advanced DMEM (Gibco) and re-suspended in ice-cold TB-ORG medium containing advanced DMEM supplemented with 1x B-27, 1x ITS-X, 10 mM HEPES (Gibco), 2 mM glutamine (Gibco), 1 µM A83-01, 100 ng/ml rhEGF and 3 µM CHIR99021. Growth-factor reduced matrigel (GFR-M; Corning) was added to reach a final concentration of 60%. Drops containing 10^5^ cells per drop in 40 µl TB-ORG medium/60% GFR-M solution were placed centrally onto each well of a 24-well plate. After 1 min of incubation at 37°C the plates were flipped upside down to ensure equal spreading of the solidifying domes. After 20 min the plates were turned again and 500 µl pre-warmed TB-ORG medium was added to each well. The medium was changed every 3-5 days and organoids were allowed to form for 7 days at P0 prior to passaging at a split ratio of 1:4. For passaging, cell recovery solution (Corning) was used according to the manufacturer’s guidelines. Organoids were overlaid with ice-cold cell recovery solution and were incubated for 40 min at 4°C to depolymerize the matrigel. After a washing step using advanced DMEM and centrifugation for 5 min at 1500 rpm, organoids were re-suspended in TB-ORG medium/60% GFR-M. Molecular characterization was performed at P1-P2. Besides, inhibition of NOTCH signalling in P1 organoids was achieved by adding either 50 µM [(2R,4R,5S)-2-benzyl-5-(Boc-amino)-4-hydroxy-6-phenyl-hexanoyl]-Leu-Phe-NH_2_ trifluoroacetate salt (L-685,458; Bachem) or 10 µM Deshydroxy LY-411575 (DBZ; Sigma-Aldrich) for 7 days. Treated organoids were fixed and subjected to immunofluorescence staining or were harvested as cell pellets for Western Blot analyses.

### Genetic Modification of TSCs

To generate the inducible DN-MAML1 expression construct, a truncated version of MAML1, fused at its C-terminus to eGFP (synthesized by GenScript), was cloned into piggyBac-Tre-Dest-rtTA-HSV-neo plasmid. Truncated MAML1 contained the amino acids 13-74, thereby only encoding the basic domain needed for interaction with NOTCH-ICD. For genetic modification, three different TSC isolations (at P3-P5) derived from different first trimester placentae were used. 1 µg piggyBac plasmid together with 1 µg transposase plasmid was transfected into 70-80% confluent TSCs using DNAfectin Plus (abm). After transfection, TSCs were selected with 250 µg/ml G418 (Gibco). After about two weeks, GFP^+^ single clones were picked and further maintained in TSC medium in the presence of 125 µg/mL G418. DN-MAML1-GFP expression was induced with 1 µg/ml doxycycline (Sigma) for 24 h up to 6 days. To generate TSCs constitutively overexpressing NOTCH3 ICD, three different TSC isolations (at P3-P5) were transfected with hNICD3(3xFLAG)-pCDF1-MCS2-EF1-copGFP (Addgene plasmid #40640; (Zhao et al., 2012)), or pcDNA3.1(-) as a control plasmid using DNAfectin Plus. Transfection rate was determined by counting GFP-positive cells. After 48 h of overexpression, cells were subjected to downstream analyses.

### Genetic Modification of Primary Cytotrophoblasts

Pooled first trimester placental tissue (6^th^-8^th^ week of gestation) was used for isolation of primary CTBs. Purification was performed as described above using three digestion steps followed by Percoll density gradient centrifugation. Before seeding onto fibronectin-coated dishes, cells were transfected with plasmids encoding hNICD3(3xFLAG)-pCDF1-MCS2-EF1-copGFP (Addgene plasmid #40640) or pcDNA3.1(-) as a control using the AMAXA SG Cell line kit (4D-Nucleofector program EO-100; Lonza). Cells were further cultivated in DMEM/F12 supplemented with 10% FBS (Sigma) and 0.05 mg/ml gentamycin for 24 h before harvesting and subjecting to downstream analyses.

### Immunofluorescence in paraffin-embedded tissues

TB-ORGs were fixed in 4% formaldehyde solution and embedded in paraffin as described elsewhere (Haider et al., 2018), while placental tissues were fixed in 7.5% formaldehyde and further processed as mentioned (Haider et al., 2016). Briefly, de-paraffinized and re-hydrated serial sections were subjected to antigen retrieval using 1x PT module buffer 1 (Thermo Scientific) for 36 min at 93°C using a KOS microwave Histostation (Milestone). Slides were then treated with blocking solution (Tris-buffered saline, pH 7.6/0.1 % Tween (TBST) supplemented with 5% normal goat serum) for 1 h at room temperature and subsequently incubated with primary antibodies (listed in Table S1) diluted in TBST / 5% normal goat serum overnight at 4°C. The following day, sections were washed three times with TBST and then incubated with appropriate secondary antibodies (2 µg/mL in TBST / 1% bovine serum albumin (BSA)) and 1 µg/mL DAPI (Roche) for one hour at room temperature. Stained sections were analyzed by fluorescence microscopy (Olympus BX50) and digitally photographed (CellP software; Olympus). Images were further processed using Adobe Photoshop CC 2023 including the automated merge function. For calculating the percentage of Ki67-positive cells in TB-ORGs, the total number of DAPI-positive nuclei and Ki67-stained nuclei were counted manually using Adobe Photoshop CC 2023.

### Immunofluorescence in cultured trophoblast stem cells

TSCs were fixed with 4% paraformaldehyde for 10 minutes at room temperature. Subsequently, cells were washed with PBS, treated with 0.1% Triton X-100/PBS for 5 min and then washed again. After a 30 min blocking step using 0.05% Fish Skin Gelatine/PBS, cells were incubated with primary antibodies (as listed in Table S1) diluted in 0.05% Fish Skin Gelatine/PBS overnight at 4°C. The following day, cells were washed with PBS and were incubated with the appropriate secondary antibody (2 µg/mL in 0.05% Fish Skin Gelatine/PBS) and 0.5 µg/mL DAPI for 1 h at room temperature. Stained cells were analyzed by fluorescence microscopy and digital pictures were taken with the EVOS FL Cell Imaging System (Life technologies). Images were further processed using Adobe Photoshop CC 2023 including the automated merge function. For quantification of cell fusion, TSC cultures were stained with syndecan-1 (SDC-1)/DAPI. ImageJ 1.52p (Wayne Rasband, National Institutes of Health, USA) was used for semi-automated quantification of SDC1^+^ areas. Thereby, stained areas were encircled manually and the measured area was calculated in mm^2^ by using the length of the incorporated scale bar to set the correct distance in pixels. The SDC1-positive area was normalized to the total number of nuclei, determined by DAPI staining.

### Proliferation Assay

DN-MAML1 transfected TSCs were cultured for 6 days with and without doxycycline stimulation prior to the addition of 10 µM 5-ethynyl-2’-deoxyuridine (EdU) (EdU-Click 488; BaseClick) overnight. Besides, NOTCH3-ICD or pcDNA3.1(-) transfected TSCs were cultured for 48 h before incubation with 10 µM EdU for 5 h. Subsequently, TSCs were fixed and EdU was detected according to the manufacturer’s instructions. In addition, nuclei were stained with 0.5 µg/ml DAPI. Cells were analyzed by fluorescence microscopy and digital pictures were taken with the EVOS FL Cell Imaging System (Life technologies). Images were processed and the percentage of proliferating cells was calculated by manually counting the numbers of total nuclei and EdU-positive nuclei using Adobe Photoshop CC 2023.

### Real-time quantitative PCR (qPCR)

RNA Isolation (PeqGold Trifst; PeqLab) and reverse transcription (RevertAid H Minus Reverse Transcriptase; Thermo Scientific) were performed according to the manufacturer’s instructions. For qPCR, the following TaqMan Gene Expression Assays (ABI) were utilized: *CCNA2* (Hs00996788_m1), *CCND1* (Hs00277039_m1), *CGB3* (Hs00361224_gH), *DLL1* (Hs00194509_m1), *DLL3* (Hs01085096_m1), *DLL4* (Hs00184092_m1), *ENDOU* (Hs00195731_m1), *GDF15* (Hs00171132_m1), *JAG1* (Hs01070032_m1), *JAG2* (Hs00171432_m1), *MAML1* (Hs00207373_m1), *MAML2* (Hs00418423_m1), *MAML3* (Hs07289055_g1), *MYC* (Hs00153408_m1), *NOTCH1* (Hs01062014_m1), *NOTCH3* (Hs01128541_m1), *NOTCH4* (Hs00965889_m1), *TEAD4* (Hs01125032_m1), *TP63* (Hs00978340_m1), *YAP1* (Hs00902712_g1). Signals (ΔCt) were normalized to TATA-box binding protein (*TBP*, 4333769F). To determine the amount of DN-MAML1 expression in TSC clones cultured in the absence or presence of doxycycline, SYBR Green dye-based qPCR detection was performed. The primers used for DN-MAML1 verification were: 5’-AGCACATGGTGAGCAAGG-3’ and 5’-GCAGATGAACTTCAGGGTCAG-3’. BrightGreen Express 2X qPCR MasterMix – Low ROX (abm) was used together with a final primer concentration of 250 nM, respectively. Cycling settings were applied according to the manufacturer’s instructions. Signals (ΔCt) were normalized to GAPDH expression (GAPDHfw70 5’-CCACCCATGGCAAATCC-3’ and GAPDHrev70 5’-GATGGGATTTGCATTGATGACA-3’; Sigma). The amplification efficiencies were determined by serial dilution and calculated as E = 10^-1/m^ × 100, where E is the amplification efficiency and m is the slope of the dilution curve. The Pfaffl method was used for calculation of the gene expression ratio (Pfaffl, 2001).

### Western Blotting

Whole-cell lysates were prepared using standard protocols as previously done (Haider et al., 2022; Meinhardt et al., 2020). Protein extracts were separated on SDS/PAA gels, transferred onto Hybond-P PVDF membranes (GE Healthcare) and incubated with primary antibodies (listed in Tables S1) overnight at 4°C. On the following day, membranes were washed and incubated for 1 h with appropriate HRP-conjugated secondary antibodies. Signals were developed using WesternBright Chemiluminescence Substrat Quantum (Biozym) and visualized with a ChemiDoc Imaging System (Biorad).

### Immunoprecipitation

Whole-cell lysates were prepared using cell lysis buffer and sonication as described in standard protocols (Cell Signaling). Immunoprecipitations were performed using NOTCH3 or MAML1 antibodies, as well as appropriate rabbit IgG controls (listed in Table S1) according to manufacturer’s instructions (Cell Signaling, no. 73778). Briefly, protein lysates were pre-cleared and then incubated with primary antibodies or IgG controls overnight at 4°C. Immuno-complexes formed between proteins and antibodies in lysates were incubated with protein A-coupled magnetic beads to form pellets. Pellets were washed with lysis buffer and then re-suspended in 3x SDS sample buffer and subjected to Western blotting analyses for the detection of co-immunoprecipitated proteins.

### Bulk RNA-Seq

For bulk RNA-seq, total RNA was prepared by using an AllPrep DNA/RNA/miRNA Universal Kit (Qiagen). Sequencing libraries were prepared at the Core Facility Genomics of the Medical University of Vienna, using the NEBNext Poly(A) mRNA Magnetic Isolation Module and the NEBNext UltraTM II Directional RNA Library Prep Kit for Illumina according to the manufacturer’s instructions (New England Biolabs). Libraries were QC-checked on a Bioanalyzer 2100 (Agilent) using a High Sensitivity DNA Kit for correct insert size and quantified using Qubit dsDNA HS Assay (Invitrogen). Pooled libraries were sequenced on two flowcells of a NextSeq500 instrument (Illumina) in 1×75bp single-end sequencing mode. On average, 33 million reads per sample were generated. FASTQ files were generated using the Illumina bcl2fastq command line tool (v2.19.1.403) including trimming of the sequencing adapters. Read quality was assessed by FASTQC.

### RNA-Seq Data Analysis

The DESeq2 package was used to analyze the count matrix, with the model being constructed based on conditions and donor id (Love et al., 2014). We applied variance stabilizing transformation for the PCA plot, followed by plotting of the key components, PC1 and PC2. The log fold change shrinkage of the results was executed using approximate posterior estimation for GLM coefficients (Zhu et al., 2019). Differentially expressed genes were identified based on an adjusted p-value less than 0.05 and an absolute log2 fold change greater than 0.5. The identified differentially expressed genes were utilized in crafting the Venn diagrams. Heatmaps were produced using the pheatmap package, with genes selected manually. The EnhancedVolcano package was used to generate volcano plots and key transcripts were labeled (Blighe K, 2023). The entire analysis was conducted using R version 4.0.4 (Team, 2022).

### Statistics

All statistical analyses were performed using GraphPad Prism 9.5. D’Agostino-Pearson normality test was performed to test Gaussian distribution, and equality of variances was examined with Bartlett’s test.

### Data Availability Statement

Raw RNA-seq data are accessible at the Gene Expression Omnibus (GEO) database (accession number GSE233140).

## Acknowledgements

The authors thank C. Fiala, Gynmed Clinic for providing placental material and for gathering written informed consent of patients. We are grateful to Core Facilities of the Medical University of Vienna, a member of VLSI, for perfoming RNA-seq. We thank P. Latos, Medical University of Vienna, for providing the piggyBac-Tre-Dest-rtTA-HSV-neo plasmid. The study was supported by the Austrian Science Funds (Project P31470-B30 to M.K and Projects P34588-B and P36159-B to S.H.).

## Author contributions

Conceptualization: S.H., B-K.K, M.K.; Methodology: B.D., A.I.L, V.K., G.M; Investigation: B.D., A.I.L, V.K., G.M; Data analyses: B.D., A.I.L, V.K., G.M; Writing, review & editing: J.P., B-K.K, S.H., M.K.; Supervision: S.H., M.K.; Project administration: M.K.; Funding acquisition: S.H., M.K.;

## Summary statement

NOTCH3 signalling has a crucial role in early development of the human placenta by promoting expansion of its founder cell, the epithelial trophoblast stem cell.

